# Detailed metabolic phenotyping of four tissue specific Cas9 transgenic mouse lines

**DOI:** 10.1101/2021.03.31.436695

**Authors:** Simon T. Bond, Aowen Zhuang, Christine Yang, Eleanor A.M. Gould, Tim Sikora, Yingying Liu, Ying Fu, Kevin I. Watt, Yanie Tan, Helen Kiriazis, Graeme I. Lancaster, Paul Gregorevic, Darren C. Henstridge, Julie R. McMullen, Peter J. Meikle, Anna C. Calkin, Brian G. Drew

## Abstract

CRISPR/Cas9 technology has revolutionized gene editing and fast tracked our capacity to manipulate genes of interest for the benefit of both research and therapeutic applications. Whilst many advances have, and continue to be made in this area, perhaps the most utilized technology to date has been the generation of knockout cells, tissues and animals by taking advantage of Cas9 function to promote indels in precise locations in the genome. Whilst the advantages of this technology are many fold, some questions still remain regarding the effects that long term expression of foreign proteins such as Cas9, have on mammalian cell function. Several studies have proposed that chronic overexpression of Cas9, with or without its accompanying guide RNAs, may have deleterious effects on cell function and health. This is of particular concern when applying this technology in vivo, where chronic expression of Cas9 in tissues of interest may promote disease-like phenotypes and thus confound the investigation of the effects of the gene of interest. Although these concerns remain valid, no study to our knowledge has yet to demonstrate this directly. Thus, in this study we used the lox-stop-lox (LSL) spCas9 ROSA26 transgenic (Tg) mouse line to generate four tissue-specific Cas9-Tg models with expression in the heart, liver, skeletal muscle and adipose tissue. We performed comprehensive phenotyping of these mice up to 20-weeks of age and subsequently performed molecular analysis of their organs. We demonstrated that Cas9 expression in these tissues had no detrimental effect on whole body health of the animals, nor did it induce any tissue-specific effects on energy metabolism, liver health, inflammation, fibrosis, heart function or muscle mass. Thus, our data suggests that these models are suitable for studying the tissue specific effects of gene deletion using the LSL-Cas9-Tg model, and that phenotypes observed utilizing these models can be confidently interpreted as being gene specific, and not confounded by the chronic overexpression of Cas9.

## Introduction

Since the discovery and proven utility of CRISPR/Cas9 based gene editing technologies, there has been a proliferation of applications that take advantage of this ground-breaking technology. Whilst the potential for this relatively simple but precise, genetic manipulation tool is obvious, the speed at which the field is developing often means that subtle off-target and deleterious effects of such an approach can be overlooked. Studies over the past 5 years have demonstrated that each system requires important optimization to ensure accurate gene editing, whilst minimizing off-target editing and potential toxicity induced by the introduction of foreign genetic machinery (Broeders et al., 2020; Molla and Yang, 2019).

A major advantage of CRISPR based editing in the pre-clinical biomedical arena is the rapid development of animal models that harbor global gene deletions or conditional targeting of alleles. These models historically took 2-3 years to generate, where now a global deletion model using CRISPR can take 3 months or less to generate (Singh et al., 2015). Moreover, CRISPR overcomes the need to generate one mouse model per gene of interest, as is the case with floxed alleles. Indeed, by over-expressing Cas9 globally in mice, or in a tissue specific manner, one can generate a model where almost any gene can be manipulated simply by introducing a guide RNA that targets your gene of interest. This flexibility of manipulation has made it feasible to use one mouse model, or even an existing disease model, to study the effect of manipulating one or many genes in combination.

One such model developed is the lox-STOP-lox spCas9-transgenic (LSL-spCas9Tg) mouse (Platt et al., 2014). This model harbors the spCas9 gene at the ROSA26 locus, but is silenced in the basal state by commonly applied repressor elements. By flanking the repressor or “STOP” element with loxP sites, the construct becomes inducible in the presence of Cre-recombinase. This model has been used by a number of groups to demonstrate robust Cas9 expression induced by Cre-recombinase, and subsequent CRISPR mediated gene editing upon the administration of a guide RNA (Laidlaw et al., 2020; Shamsi et al., 2020; Zhu et al., 2020).

One concern with such an approach is whether chronic expression of the foreign CRISPR machinery mediates phenotypic effects *per se* (Broeders et al., 2020). To this end, we have successfully developed four tissue-specific Cas9 transgenic mouse models that include key tissues of interest in the metabolism field (skeletal muscle, liver, adipose tissue and heart). We phenotyped these animals for readouts of whole body metabolism, adiposity, glucose tolerance, toxicity and cardiac function, and demonstrated that none of the models were impacted by the chronic (~12 weeks) presence of Cas9. These findings provide important and much needed confidence for researchers who wish to use these mouse models in the future, who can now confidently ascribe any observed metabolic phenotype to their gene of interest, and not to underlying unwanted side effects of the model itself.

## Results

### Tissue Specific and Inducible Expression of Cas9 in four mouse models

To generate tissue specific Cas9 transgenic mice, we crossed the LSL-spCas9 Tg mouse (Platt et al., 2014) with four different tissue specific Cre-recombinase transgenic mouse lines. Three of these models (ACTA1-Cre, AdipoQ-Cre and MHCalpha-Cre) were tamoxifen inducible via the use of Cre-recombinase that was fused to the modified estrogen receptor (mER), often referred to as ERT2. The other model (albumin-Cre) was constitutively active and expressed from the albumin promoter (Alb-Cre). These breeding strategies resulted in four separate mouse models with an expected tissue specific expression of Cas9 in liver (Albumin), heart (MHC-alpha), skeletal muscle (ACTA1), and adipose tissue (AdipoQ), (**Figure 1A**).

**Figure 1:**
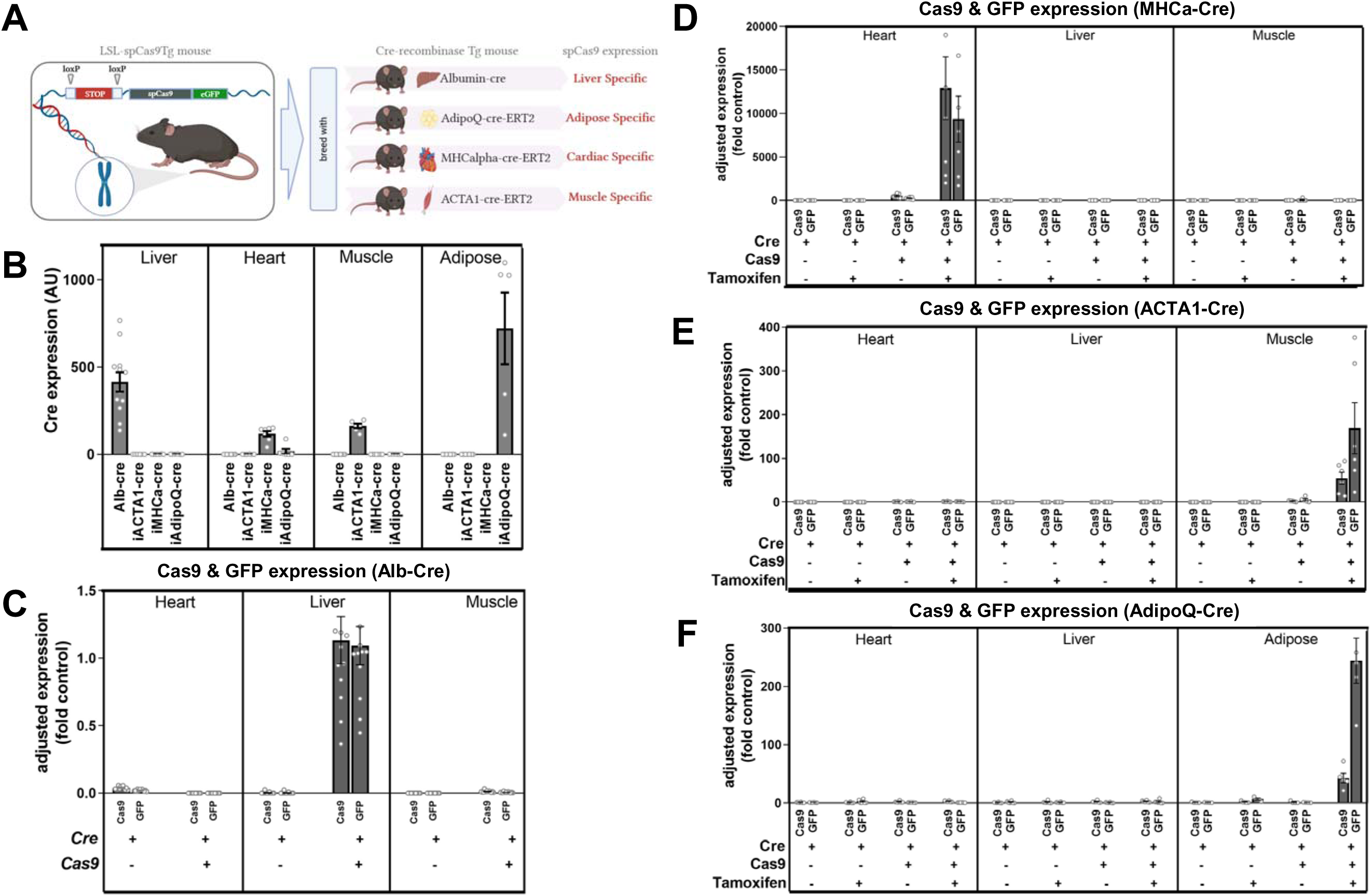
Tissue Specific and Inducible Expression of Cas9 in four mouse models. **A.** Schematic outlining the breeding strategy and generation of the four tissue specific Cas9 transgenic mouse lines. The LSL-spCas9Tg mouse was bred with four different Cre-lines (Albumin-Cre = Liver, AdipoQ-Cre-ERT2 = adipose, MHC-alpha-Cre-ERT2 = cardiac, ACTA1-Cre/-ERT2 = skeletal muscle) to generate lines that were independently maintained and studied. **B.** Relative Cre-recombinase expression as determined by qPCR for each line (annotated across the bottom) in the various tissues (annotated across the top) for each mouse line. Data are normalized to a housekeeping gene (*Rplp0*) and presented as arbitrary unit (AU) for “Cas9+Cre Active” compared to “Cre active” groups. Adjusted expression of Cas9 and GFP as determined by qPCR and presented as fold change to control in the heart, liver and muscle of **C.** spCas9Tg+Alb-Cre constitutively active Cre line, showing just the two groups per tissue = Cre-Cas9 and Cre+Cas9 as indicated by the (+) and (-) signs at the bottom of the graph, Cas9 and GFP in the **D.** spCas9Tg+MHC-alpha-Cre/-ERT2 mice, **E.** spCas9Tg+ACTA1-Cre/-ERT2 mice and **F.** spCas9Tg+AdipoQ-Cre/-ERT2 mice. Because the MHC-alpha-, ACTA1- and AdipoQ-Cre are inducible (tamoxifen, TAM) lines, there are four groups per tissue = Cre-Cas9 (no TAM), Cre-Cas9 (plus TAM), Cre+Cas9 (no TAM) and Cre+Cas9 (plus TAM) as indicated by the (+) and (-) signs at the bottom of the graphs. All data are presented as mean±SEM, n=4-12/group. LSL = lox-STOP-lox, spCas9 = Cas9 from *S. Pyogenes*, Tg = transgenic, eGFP = enhanced green fluorescent protein, ERT2 = 2 x tamoxifen sensitive mutant estrogen receptor

Because three of the Cre-models were inducible (with tamoxifen), we administered tamoxifen or vehicle (oil) using specific regimens for each model (see methods) between 6-8 weeks of age, then allowed 2 weeks for maximal gene expression before any phenotyping was performed. For the inducible models, we studied four cohorts of mice per model, which included: Cre+OIL (Cre-inactive), Cas9+Cre+OIL (Cas9+Cre-inactive), Cre+Tamoxifen (Cre-active) and Cas9+Cre+Tamoxifen (Cas9+Cre-active) mice. For the constitutively active Albumin-Cre model, there was no oil/tamoxifen treatment so there were only two cohorts; Cre-active and Cas9+Cre-active.

To characterize the different models, we performed phenotyping that included assessment of body weight, fat mass and lean mass by EchoMRI, and glucose tolerance tests (GTTs) which were performed at various times for different models over the subsequent 12 weeks. We also performed some model specific phenotyping, including echocardiography of the MHC-alpha model to analyze heart function.

At the completion of each study (mice up to approximately 20-22 weeks of age), tissues were collected, processed and analyzed to first confirm that each model displayed the expected expression profiles. Using qPCR analysis, we demonstrated that each model exhibited tissue specific Cre-recombinase expression, with robust expression in the expected tissue, but no detected expression in other tissues (**Figure 1B**). Importantly, we also demonstrate using qPCR that Cas9 expression was tissue specific, and that this expression was dependent on both Cre-expression and the administration of tamoxifen in the inducible models. As a secondary confirmation we also determined the abundance of GFP by qPCR, which is co-expressed from the same transgene cassette as Cas9, but is independently processed by the ribosome (i.e. not tagged to Cas9). We demonstrated that both Cas9 and GFP exhibited the expected tissue specific expression profiles including liver (**Figure 1C)**, heart (**Figure 1D**), muscle (**Figure 1E**) and white adipose tissue (WAT, **Figure 1F**). The level of Cas9 induction varied slightly across the lines, which is likely a reflection of local transcriptional machinery, number of cells per unit of tissue, and the differential activity of Cre-recombinase in each tissue.

### Tissue Specific Expression of Cre-Recombinase, Cas9 and GFP does not Impact Animal Body Weight or Tissue Weights

Upon demonstrating that each model expressed Cas9 in a tissue specific manner, we next sought to test previously raised concerns that chronic over-expression of “foreign” enzymes such as Cre and Cas9 in metabolic tissues, might lead to phenotypic differences in animal growth and development. A simple way of testing for toxicity or growth inhibition is to compare body weight and individual tissue weights from each model at study end. We demonstrate that body weights for each model were comparable between cohorts at the time of cull, with no significant differences in body weight, whether they were expressing Cre, Cas9, or if they had been treated with tamoxifen/oil (**Figure 2A**). Moreover, assessment of liver, WAT, Muscle (*Tibialis anterior*; TA) and heart weights at the time of cull, demonstrated no difference in the weight of any of these tissue between the various groups within each model (**Figures 2B–2E**).

**Figure 2:**
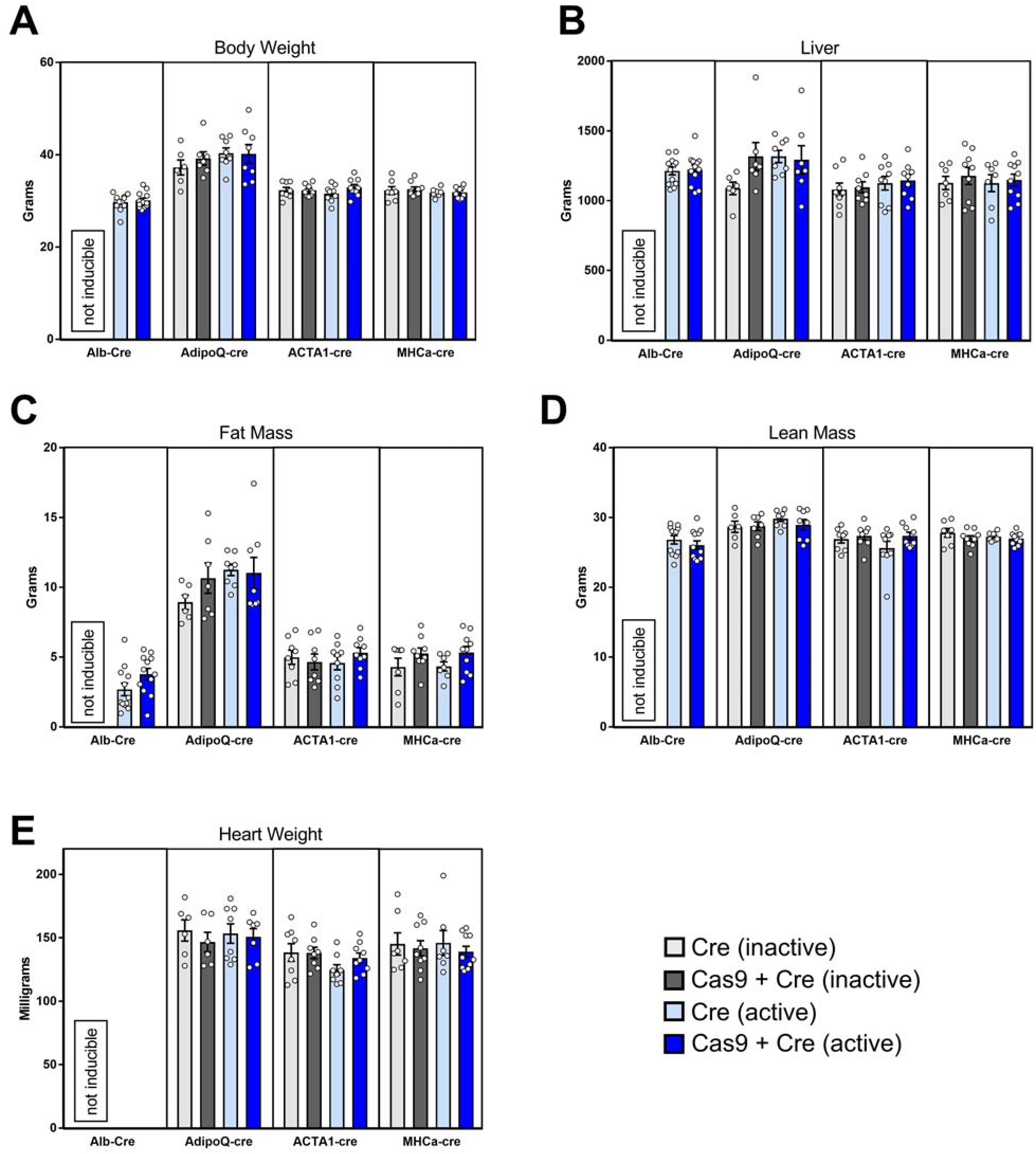
Tissue Specific Expression of Cre-Recombinase, Cas9 and GFP does not Impact Animal Body Weight or Tissue Weights. All four tissue specific Cas9 mouse lines were aged to approximately 18-20 weeks of age and analyzed for **A.** Body weight, **B.** Liver weight at cull, **C.** Fat mass by EchoMRI (final two weeks), **D.** Lean mass by EchoMRI (final two weeks) and **E.** heart weight at cull. The four groups per line are as follows: Cre (inactive – light grey), Cre+Cas9 (inactive – dark grey), Cre (active – light blue) and Cre+Cas9 (active – dark blue). All data are presented as mean÷SEM, n=6-11/group. Albumin Cre-line is constitutively active so the inactive groups were designated as “not inducible”. ND = not determined in this line.

### Tissue Specific Expression of Cre-Recombinase, Cas9 and GFP does not Alter Molecular or Physiological Readouts of Tissue Function

Whilst it was important to demonstrate that there was no effect of Cas9 expression on gross tissue weights or animal growth, we also sought to investigate whether tissue specific pathways were being impacted by chronic over expression of Cas9 in each tissue. Therefore, we performed a series of analyzes on each tissue to investigate these parameters, utilizing qPCR, histology and functional assessment.

In the liver specific Cas9 model (Alb-Cre), we used qPCR to analyze the expression of genes that were representative of pathways that provided insight into the health and activity of the liver. These included *Col1a2* and *Vim* as markers of fibrosis, *Chop* for ER stress, *Plin2* for lipid storage and *Tnfa* and *Il1b* for inflammation (**Figure 3A**). We demonstrated that none of these genes were differentially expressed in mice with livers expressing Cas9 compared to control livers, indicating that the expression of Cas9 in the liver was not impacting on liver inflammation, fibrosis or lipid handling. Moreover, representative histological sections stained with hematoxylin and eosin (H&E) demonstrated no gross changes in liver morphology (**Figure 3B**).

**Figure 3:**
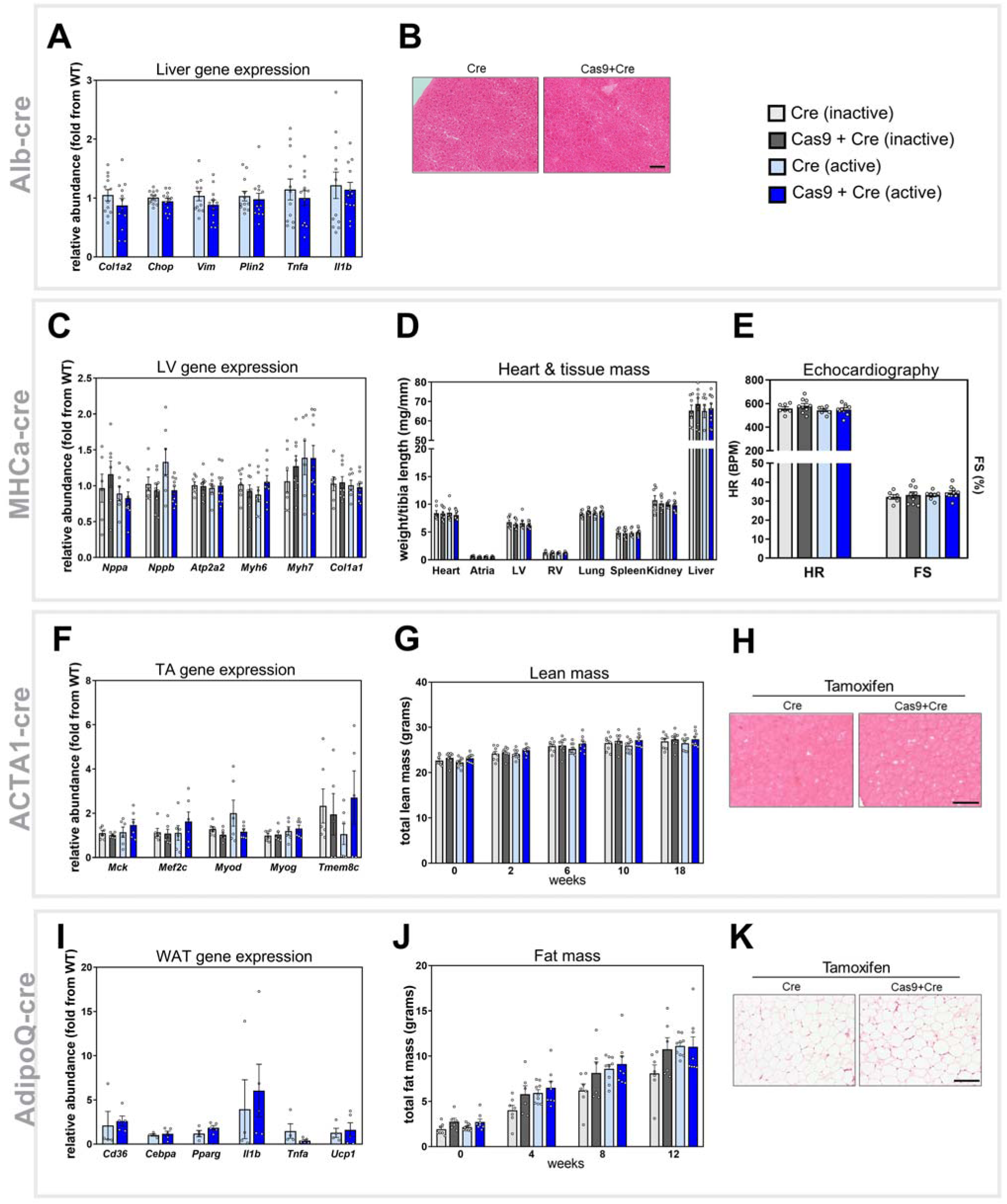
Tissue Specific Expression of Cre-Recombinase, Cas9 and GFP does not Alter Molecular or Physiological Readouts of Tissue Function. Molecular and functional read outs of tissue function specific were performed. Alb-Cre line (liver specific) was investigated for changes in **A.** hepatic mRNA expression for pathways indicative of fibrosis (*Col1a2, Vim*), ER stress (*Chop*), lipid metabolism (*Plin2*) and inflammation (*Tnfa, Il1b*) as analyzed by qPCR (n=12/group), and **B.** representative images of liver tissue morphology as assessed by histology with H&E staining. The MHC-alpha-Cre-ERT2 line (cardiac specific) was analyzed for changes in **C.** cardiac mRNA expression for pathways indicative of cardiac pathology (*Nppa, Nppb, Atp2a2, Myh6* and *Myh7*) and fibrosis (*Col1a1*), **D.** Mass of tissues pertinent to cardiac pathology (heart, atria, left ventricle (LV), right ventricle (RV), and lung) and whole body animal health (spleen, kidney and liver) and **E.** measurement of heart function including heart rate and fractional shortening (FS%) as analyzed by echocardiography, n=6-10/group. The ACTA1-Cre-ERT2 line (muscle specific) was investigated for changes in **F.***Tibialis anterior* (TA) mRNA expression for pathways indicative of muscle maturation (*Mck*), regeneration (*Mef2c, Myod* and *Myog*) and fusion (*Tmem8c*) as analyzed by qPCR, **G.** Temporal changes in lean (muscle) mass during the study (at timepoints indicated on graph) as analyzed by EchoMRI and **H.** muscle (TA) morphology as assessed by histology with H&E staining n=5-8/group. The AdipoQ-Cre/ERT2 line (adipose specific) was investigated for changes in **F.** white adipose tissue (WAT) gene expression for pathways indicative of lipid uptake (*Cd36*), adipocyte differentiation (*Cebpa, Pparg*) inflammation (*Tnfa, Il1b*) and WAT browning (*Ucp1*) as analyzed by qPCR, **G.** Temporal changes in fat (adipose) mass during the study (at timepoints indicated on graph) as analyzed by EchoMRI and **H.** WAT morphology as analyzed by histology with H&E staining, n=4-9/group. All data are presented as mean±SEM, scale bar represents 100μm.

In the heart specific Cas9 model (MHCa-Cre), we used qPCR to measure pathways in the heart that are indicative of cardiac health and function. These included classic molecular markers of heart failure including Atrial Natriuretic Peptide (ANP) (*Nppa*) and B-type Natriuretic Peptide (BNP) (*Nppb*) as well as αMHC (*Myh6*) and ßMHC (*Myh7*). We also measured a marker of fibrosis (*Col1a1*) and a marker of cardiac contractile function via the sarcoplasmic reticulum /endoplasmic reticulum Ca2+ ATPase 2a (SERCA; *tp2a2*) (**Figure 3C**). We demonstrate that were no differences in the expression of any of these markers in the left ventricle (LV) across the four groups of mice, implying that these pathways were not altered by the expression of Cas9 (or Cre-recombinase). This was supported by data demonstrating that the weight of the whole heart and the different regions of the heart from these mice including; atria, LV and right ventricle (RV), were also not different between groups (**Figure 3D**). Consistently, the lung weights, spleen weight and kidney weight (which are useful readouts of health in cardiac models) were all comparable across groups demonstrating no peripheral effects of cardiac specific overexpression of Cas9. Lastly, we assessed heart function in these mice using echocardiography. We demonstrated that at comparable heart rates (HR), under anesthesia there were no differences in fractional shortening (FS%) – a measure of systolic function, between the four groups (**Figure 3E**). Thus, collectively these data demonstrate that overexpression of Cas9 in cardiomyocytes has no impact on heart health and function.

Specific phenotyping of the muscle specific Cas9 model (ACTA1-Cre) was also performed using qPCR. We measured the expression of muscle specific genes that are known readouts of muscle development and growth in the TA muscle. These included myogenic transcription factors *Myod, Myog* and *Mef2c*, as well as the pro-fusion protein myomaker (*Tmem8c*) and mature muscle marker *Mck* (**Figure 3F**). As with our previous models, we demonstrated no difference in the expression of these genes between the four groups of mice, indicating that there were no major differences in the growth and function of adult skeletal muscle in the presence of Cas9 expression. In support of this data, using EchoMRI we demonstrated that there was no difference in lean muscle mass across the four groups, at any time point throughout the study period (**Figure 3G**). Finally, histological analyzes of TA muscle sections using H&E staining, indicated that there were no major morphological differences in the muscle structure between Cas9 positive and Cas9 negative mice (**Figure 3H**). Collectively, these data indicate that Cas9 expression in skeletal muscle has no impact on muscle health and maturation.

The final model we characterized was the adipose-specific Cas9 mouse (AdipoQ-Cre). As with previous models, using qPCR we demonstrated that there was no major difference in the expression of genes related to adipocyte differentiation and health in WAT, including the adipogenic transcription factors PPARgamma (*Pparg*) and C/EBPalpha (*Cebpa*), the lipid transporter *Cd36*, inflammatory markers *Tnfa* and *Il1b*, and the browning marker *Ucp1* (**Figure 3I**). Moreover, using EchoMRI we demonstrated that there was no difference in fat mass between the four groups in this AdipoQ-Cre model at any time point throughout the study period (**Figure 3J**). Finally, histological assessment of WAT sections using H&E staining, indicated that there were no major morphological differences in adipocyte size or structure between Cas9 positive and Cas9 negative mice (**Figure 3K**). Collectively, these data indicate that Cas9 expression in adipose tissue has no impact on WAT health and development.

### Tissue Specific Expression of Cre-Recombinase, Cas9 and GFP does not Alter Whole Body Glucose Homeostasis

Given that many groups which study metabolism have an interest in glucose homeostasis and how it relates to tissues such as liver, adipose, skeletal muscle and the heart, we sought to determine if the expression of Cas9 in these tissue led to any changes in whole body glucose handling. To investigate this, we performed fasting blood glucose measurements and oral glucose tolerance tests on all groups and models within the final two weeks of the study period. We demonstrated that there was no difference in fasting blood glucose levels between Cas9 positive and Cas9 negative mice in each of the four tissue specific mouse models (**Figures 4A, 4D, 4G and 4J**). In order to test the glucose tolerance of these models, we challenged each cohort with a standardized oral dose of glucose (2mg/kg lean mass), and subsequently measured their blood glucose concentration over two hours in an oral glucose tolerance test (oGTT). We demonstrated that all groups and models showed a peak glucose concentration of approximately 18-20mmol/L at 15 minutes post glucose delivery, which mostly returned to baseline by 60 minutes after delivery of the glucose bolus (**Figures 4B, 4E, 4H and 4K**). We also demonstrated that there was no difference in the clearance of glucose across any of the groups and in each of the models, indicating that there was no difference in the glucose tolerance of these animals. This is further demonstrated quantitatively by assessing the 90 minute cumulative area under the curve (AUC) for the tolerance test (**Figures 4C, 4F, 4I and 4L**), confirming that there was no difference in glucose tolerance between the groups in each tissue specific Cas9 model.

**Figure 4:**
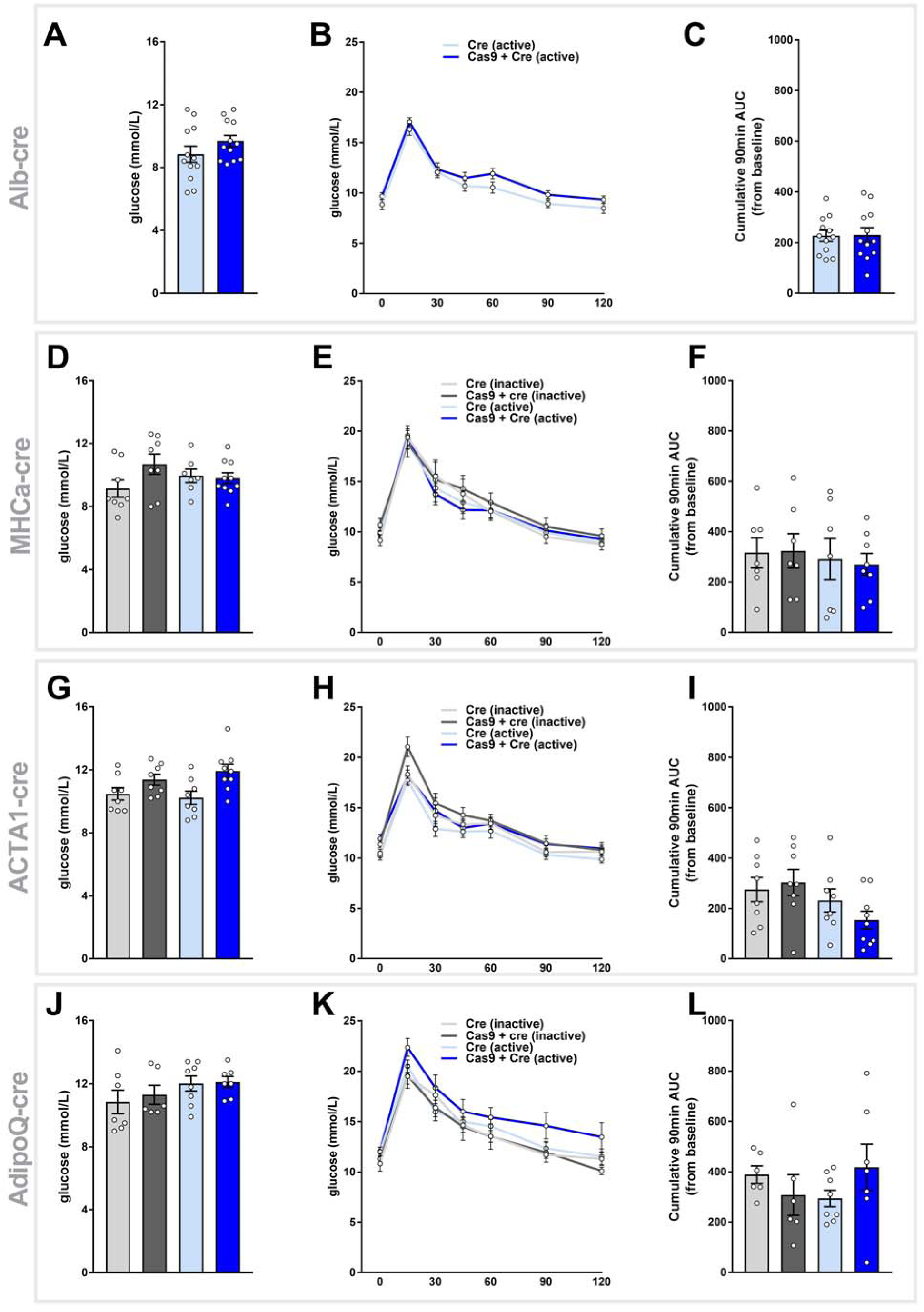
Tissue Specific Expression of Cre-Recombinase, Cas9 and GFP does not Alter Whole Body Glucose Homeostasis. All four Cas9 lines were phenotyped for parameters of whole body glucose homeostasis at the end of the study. This included assessment of fasting blood glucose for **A.** Alb-Cre (n=11-12/group), **D.** MHC-alpha-Cre-ERT2 (n=7-10/group), **G.** ACTA1-Cre-ERT2 (n=8-9/group) and **J.** AdipoQ-Cre-ERT2 (n=6-8/group) and two-hour glucose tolerance as performed by oral glucose tolerance tests (oGTT) on **B.** Alb-Cre **E.** MHC-alpha-Cre-ERT2 **H.** ACTA1-Cre-ERT2 and **K.** AdipoQ-Cre-ERT2 mice as quantified by 90 minute cumulative area under the curve (AUC) analysis for **C.** Alb-Cre **F.** MHC-alpha-Cre-ERT2 **I.** ACTA1-Cre-ERT2 and **L.** AdipoQ-Cre-ERT2. All data are presented as mean±SEM.

Collectively, the data presented above demonstrates that long term overexpression of Cas9 in four different tissue specific models, does not lead to any effects on body weight, tissue weight, the expression of markers of pathological pathways, or readouts of whole body glucose metabolism. These findings provide an important foundation for future studies that wish to use the LSL-Cas9 mouse model to study their gene of interest, and affords confidence to researchers that metabolic phenotypes they measure are unlikely to be impacted by the chronic over expression of Cas9 or Cre-recombinase in these models.

## Discussion

The discovery and implementation of CRISPR Cas9 as a gene editing tool has far reaching implications for furthering knowledge gain in biomedical science. The flexibility and comparatively simple execution of this technology means it can be utilized by most research laboratories around the world, accelerating the opportunity for discovery by several fold over existing technologies. Whilst CRISPR Cas9 has indeed been adopted quickly and efficiently by the scientific community, this has often been accomplished without due consideration for the potential negative effects on metabolic readouts that might arise from such methodologies, particularly if appropriate optimisation has not been performed. Many studies have shown that spurious gene editing can occur in the setting of chronic, high level expression of Cas9 (Hendel et al., 2015) and their accompanying single guide RNAs (sgRNAs) (Fu et al., 2013; Link et al., 2018), whilst others have expressed concern over the impacts of long term exogenous expression of CRISPR machinery (Cas9/sgRNAs), causing unwanted effects on target cells (Charlesworth et al., 2019; Enache et al., 2020). Such unwanted effects on targets cells would be particularly concerning in the in vivo setting, where even minor disruptions to tissue function over many months has the potential to substantially impact on animal health and disease risk. Unfortunately, the impact of these unwanted effects is mostly unknown at this point, and is difficult to predict without performing the experiments directly. This is likely time consuming and laborious, and thus these important “control group comparisons” are often the first experiments to be overlooked when designing new CRISPR editing experiments.

With regard to in vivo CRISPR editing, the generation of the inducible spCas9 transgenic mouse (LSL-spCas9Tg) by Zhang and colleagues has been an important tool to enable tissue and temporal specific Cas9 expression in mice (Platt et al., 2014). This model has been used in several labs around the world to successfully delete genes of interest, most commonly in myeloid or neuronal cell lineages (Laidlaw et al., 2020; Shamsi et al., 2020; Zhu et al., 2020). Whilst there are obvious advantages to using this LSL-spCas9Tg mouse model, it is less obvious what the potential disadvantages are - if any do exist. This is particularly true when generating tissue specific Cas9 models for the first time, as there would be no data available as to whether chronic Cas9 overexpression will impact the tissue of interest and the whole body phenotypes of interest.

Given our group has a major interest in metabolism and the organs that regulate whole body energy status, we are constantly performing studies in pertinent metabolic tissues such as the liver, muscle, adipose and heart. Unfortunately, to date, few studies have performed in vivo CRISPR editing in these tissues using the LSL-spCas9Tg mouse, and thus it is unclear as to whether this model would be suitable for investigating CRISPR-mediated gene deletion in the aforementioned tissues, without the risk of unwanted side effects due to chronic Cas9 overexpression.

We thus generated liver-, muscle-, adipose- and heart-specific Cas9 expressing mice using the LSL-Cas9Tg model, and demonstrated that these four models appear to be unaffected by the chronic expression of Cas9 and/or Cre-recombinase. We performed comprehensive metabolic phenotyping of all four lines including body composition, glucose tolerance, molecular and biochmical measurements, and functional readouts on various tissues. These analyses demonstrated clear tissue specificity of Cas9 expression, driven by temporal activation of Cre-recombinase using tamoxifen in all three of the inducible lines (muscle, adipose and heart), as well as the constitutive albumin (liver) line. Importantly, we were unable to detect any deleterious effects on metabolic pathways, morphology of tissues, body composition or glucose tolerance in any lines over-expressing Cas9, supporting the notion that there is no negative impact of chronic Cas9 expression in these tissues. Moreover, echocardiography also demonstrated no impact on systolic heart function in cardiac-specific Cas9 expressing mice after 20 weeks of induction, providing evidence that heart function in these mice was not affected by chronic Cas9 expression.

In summary, we provide critical evidence that the metabolism and general health of four different metabolic tissue specific mouse lines are unaffected by the chronic expression of Cas9. These findings provide confidence for researchers moving forward, who wish to use these Cas9 mouse models to manipulate the expression of genes in these particular tissues. The minimal impact of Cas9 in these studies will likely reduce the need for future studies to perform specific controls groups, reducing animal numbers and sparing expensive resources. Our data will also provide confidence that observed phenotypes related to gene deletions in these models in future studies, are likely to be specific to the gene of interest rather than being related to the chronic over-expression of Cas9. Thus these findings provide an important resource for the research community.

## Methods

### Generation of Tissue Specific Cas9 Animal Models

All animal experiments were approved by the Alfred Research Alliance (ARA) Animal Ethics committee (E/1756/2017/B) and performed in accordance with the research guidelines of the National Health and Medical Research Council of Australia. Tissue specific expression of Cas9 was achieved using the Cre-Lox system, where Cre-recombinase was used to remove the “STOP” sequence from the lox-stop-lox (LSL) cassette separating the promoter and Cas9 genes in the mouse described by Zhang et al (Platt et al., 2014). The four different mouse lines were generated by crossing the LSL-spCas9-Tg mouse with either the Albumin-Cre, ACTA1-Cre-ERT2, AdipoQ-Cre-ERT2 or MHCalpha-Cre-ERT2 mice. All mice were on a C57BL/6J background and are available from Jackson Laboratories. We generated two cohorts of n=8-10 male mice of Alb-Cre-Cas9Tg mice (Cre & Cre+Cas9), and 2 cohorts of n=16-20 male mice of the inducible Cre-Cas9Tg mouse lines (Cre & Cre+Cas9), the latter of which were split further into two groups each and treated with either vehicle (sunflower oil) or Tamoxifen in sunflower oil, generating the following four groups; Cre inactive (OIL), Cre+Cas9 inactive (OIL), Cre active (tamoxifen) and Cre+Cas9 active (tamoxifen). Only the final group (Cre+Cas9 active) was expected to express Cas9, with the others serving as either Cre or tamoxifen control groups.

### Animal Treatments and Husbandry

All mice were bred and sourced through the ARA Precinct Animal Centre and randomly allocated into their respective groups. For tamoxifen inducible models, they were treated as follows. For ACTA1-Cre-ERT2 and AdipoQ-Cre-ERT2 models, mice were aged to 6-8 weeks old before being gavaged with either Tamoxifen (80mg/kg) in sunflower oil, or sunflower oil alone, for 3 consecutive days. For the MHC-alpha-Cre-ERT2 model, mice were IP injected once with 40mg/kg of Tamoxifen in sunflower oil, or sunflower oil alone. Following tamoxifen treatment, mice were left to recover for 2 weeks, after which they were maintained on a normal chow diet (Normal rodent chow, Specialty feeds, Australia) and housed at 22°C on a 12hr light/dark cycle with access to food and water *ad libitum* with cages changed weekly for 12 weeks. Cohorts of mice were subjected to EchoMRI and body weight analysis throughout the study period. In the last two weeks of the study period, all animals underwent oral glucose tolerance tests, whilst the MHC-alpha mice were also subjected to cardiac function assessment via echocardiography. At the end of the study, mice were fasted for 4-6 hours and then anesthetized with a lethal dose of ketamine/xylazine before blood and tissues were collected, weighed and snap frozen for subsequent analysis.

### Glucose Tolerance Tests

Oral glucose tolerance tests (oGTT) were performed as previously described (Bond et al., 2019a; Bond et al., 2021). In the final two weeks of the study period mice were fasted for 4-6 hours and gavaged at a glucose dose of 2g/kg of lean mass as determined by EchoMRI. Blood glucose was determined using a glucometer at the following times points; 0, 15, 30, 45, 60, 90 and 120 minutes.

### EchoMRI

Body composition was analyzed using the 4 in 1 NMR Body Composition Analyzer for Live Small Animals, according to the recommendations of the manufacturer (EchoMRI LLC, Houston, TX, USA). This provides measurements of lean mass and fat mass in living animals as previously described (Bond et al., 2019a; Bond et al., 2021).

### Histology

Liver and muscle were embedded cut side down in OCT before being frozen in a bath of isopentane submerged in liquid nitrogen. After freezing, blocks were brought to −20°C and 5μm sections were cut using a Leica Cryostat. Sections were mounted and dried overnight at room temperature before being fixed in Methanol. WAT samples were fixed in formalin and mounted in Paraffin, before 5μm sections were cut on a Leica microtome. All sections were stained with hematoxylin and eosin and slide images were captured using Olympus Slide scanner VS120 (Olympus, Japan) and viewed in the supplied program (OlyVIA Build 13771, Olympus, Japan).

### Quantitative PCR (qPCR)

RNA was isolated from tissues using RNAzol reagent and isopropanol precipitation as previously described (Bond et al., 2021; Bond et al., 2019b). Briefly, cDNA was generated from RNA using MMLV reverse transcriptase (Invitrogen) according to the manufacturer’s instructions. qPCR was performed on 10ng of cDNA using the SYBR-green method on an ABI 7500, using primer sets outlined in Table 1. Primers were designed to span exon-exon junctions where possible, and were tested for specificity using BLAST (Basic Local Alignment Search Tool; National Centre for Biotechnology Information). Amplification of a single amplicon was estimated from melt curve analysis, ensuring only a single peak and an expected temperature dissociation profile were observed. Quantification of a given gene was determined by the relative mRNA level compared with control using the delta-CT method, which was calculated after normalisation to the housekeeping gene *Ppia* or *Rplp0*.

**Table 1.**
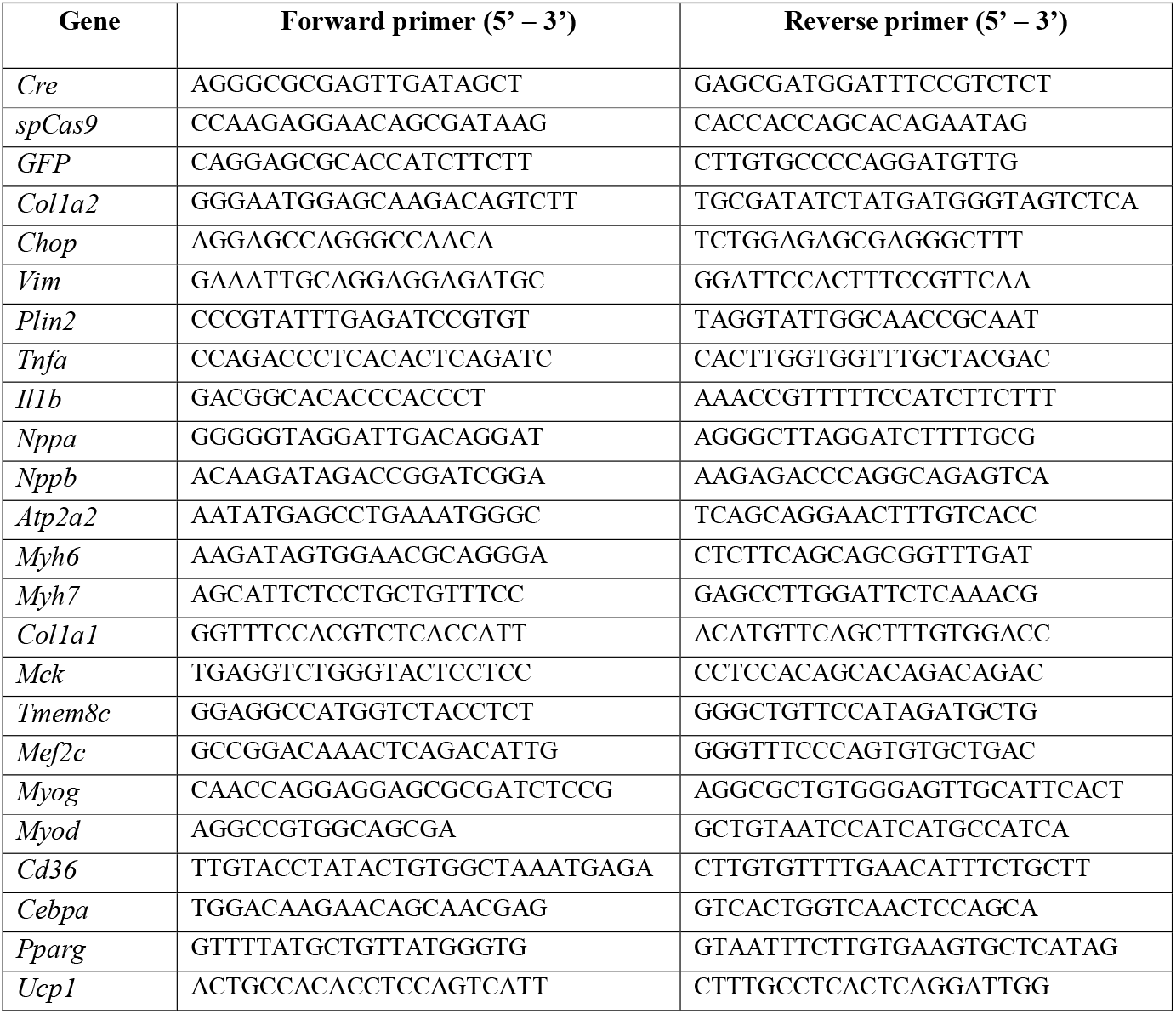
Forward and reverse primer sets for detection of the designated mouse genes using qPCR.

### Echocardiography

Echocardiography was performed on mice anaesthetised with isoflurane (1.5-2%) at the end of the 12-week period following tamoxifen induction, using a 15-MHz linear transducer L15-7io with a Philips iE33 Ultrasound Machine (North Ryde, NSW, Australia). Data were analyzed and verified by two independent researchers according to QC procedures and validation measures as outlined previously (Donner et al., 2018).

### Data Inclusion and Exclusion Criteria

For animal experiments, phenotyping data points were excluded using the following pre-determined criteria: if the animal was unwell at the time of analysis, there were identified technical issues (such as unclear signal from echocardiography) or data points were identified as outliers using Tukey’s Outlier Detection Method (Q1 minus 1.5 IQR or Q3 plus 1.5 IQR). If repeated data points from the same mouse failed QC based on pre-determined criteria, or several data points were outliers as per Tukey’s rule, the entire animal was excluded from that given analysis (i.e. during glucose tolerance tests, indicating inappropriate gavage). For in vivo and in vitro tissue and molecular analyzes, data points were only excluded if there was a technical failure (i.e. poor RNA quality, failed amplification in qPCR), or the value was biologically improbable. This was performed in a blinded fashion (i.e. on grouped datasets before genotypes were known).

## Acknowledgements

We acknowledge funding support from the Victorian State Government OIS program to Baker Heart & Diabetes Institute. These studies were supported by funding from the Baker Heine Trust through the both Obesity & Lipid Program and the Bioinformatics Programs, as well as the Baker Bertalli mini-grant scheme at Baker. We thank members of the MMA, LMCD, Cardiac Hypertrophy, Metabolomics, and Hematopoiesis & Leukocyte Biology laboratories at BHDI for their contributions. We also acknowledge the use of the facilities and technical assistance of the Monash Histology Platform, Department of Anatomy and Developmental Biology, Monash University

## Author contributions

BGD and ACC designed and conceived the study. BGD wrote the manuscript and all other authors read and/or edited the manuscript. BGD, STB, DCH, AZ, CY, EAMG, YL, HK, KIW and GIL performed animal experiments and phenotyping. BGD, STB, AZ, TS, YL, YF and YT analyzed data, processed tissue samples and performed molecular and biochemical experiments. PG, JRM, PJM and ACC provided reagents, experimental advice and access to infrastructure and resources.

